# Uterine NK cells underexpress receptors recognizing HLA-C2 and HLA-G in reproductive failure

**DOI:** 10.1101/2022.11.25.517971

**Authors:** Ee Von Woon, Dimitrios Nikolaou, Kate MacLaran, Julian Norman-Taylor, Priya Bhagwat, Antonia O. Cuff, Mark R. Johnson, Victoria Male

**Affiliations:** Department of Metabolism, Digestion and Reproduction, Institute of Developmental Reproductive and Developmental Biology, Imperial College London, Chelsea and Westminster Hospital Campus, 369 Fulham Road, London, SW10 9NH, United Kingdom; The Fertility Centre, Chelsea and Westminster Hospital, 369 Fulham Road, London, SW10 9NH, United Kingdom; Department of Cellular Pathology, Imperial College Healthcare NHS Trust, Charing Cross Hospital, Fulham Palace Road, W6 8RF United Kingdom

**Author notes:** **Correspondence:** Ee Von Woon.

**Keywords:** NK cells, endometrium, pregnancy, ILC, recurrent miscarriage, infertility, recurrent implantation failure, reproductive failure

## Abstract

A significant proportion of recurrent miscarriage, recurrent implantation failure and infertility are unexplained, and these conditions have been proposed to have an etiology of immunological dysfunction at the maternal-fetal interface. Uterine Natural Killer cells (uNK) comprise three subsets and are the most numerous immune cells found in the uterine mucosa at the time of implantation. They are thought to play an important role in successful pregnancy by regulation of extravillous trophoblast (EVT) invasion and spiral artery remodelling. Here, we examine the frequency, phenotype and function of uNK1-3 from the uterine mucosa of 16 women with unexplained reproductive failure compared to 11 controls with no reproductive problems, during the window of implantation. We report that KIR2DL1/S1 and LILRB1 expression is lower in the reproductive failure group for both uNK (total uNK, uNK 2 and 3) and pNK. We also show that degranulation activity is significantly reduced in total uNK, and that TNF-α production is lower in all uNK subsets in the reproductive failure group. Taken together, our findings suggest that reproductive failure may be caused by global reduction in expression of uNK receptors important for interaction with HLA-C and HLA-G on EVT during early pregnancy, leading to reduced uNK activation. This is the first study to examine uNK subsets during the window of implantation in women with reproductive failure and will serve as a platform to focus on particular aspects of phenotype and function of uNK subsets in future studies. Further understanding of uNK dysregulation is important to establish potential diagnostic and therapeutic targets in the population of women with unexplained reproductive failure.

## Introduction

Subfertility affects up to 1 in 7 couples and is commonly defined as failure to conceive after regular unprotected sexual intercourse for 12 months (*World Health Organization*, 2019; Carson and Kallen, 2021). Recurrent miscarriage (RM) can be defined as loss of two (American Society for Reproductive Medicine, 2012; Bender Atik *et al*., 2018), three (RCOG, 2011) or more clinical pregnancies and affects 1-2% of couples trying to conceive. Recurrent implantation failure (RIF) is a term used to describe failure to conceive after several attempts of in vitro fertilization (IVF). Similar to recurrent miscarriage, it has variable definitions which involve failure to achieve pregnancy after two (Polanski *et al*., 2014) or three (Margalioth *et al*., 2006; Coughlan *et al*., 2014; Mascarenhas *et al*., 2021) fresh or frozen cycles with transfer of good quality embryos in women under the age of 40. Although these three conditions represent different clinical entities, a significant proportion of the cases in each group remain unexplained. Indeed, 1 in every 4 cases of subfertility do not have a cause identified after routine investigations (unexplained infertility: UI) (*NICE*, 2013; Carson and Kallen, 2021) and up to 50% cases of recurrent miscarriage (Rai and Regan, 2006).

It has been suggested that these conditions may have an immune etiology, resulting from dysfunctional molecular mechanisms involving the immune system at the maternal fetal interface. Indeed, immune dysregulation has been proposed to cause defective implantation and deep placentation resulting in a spectrum of pregnancy disorders such as recurrent miscarriage, pre-eclampsia, intra-uterine growth restriction and preterm labour, collectively termed “Great Obstetrical Syndromes” (Brosens *et al*., 2011; Moffett and Shreeve, 2022). In line with this, the term “reproductive failure” was coined based on the theory that RM and RIF have a common pathogenic pathway involving a complex interplay of hormonal and immunological factors to ensure optimum endometrial receptivity (Makrigiannakis *et al*., 2011). Here, we use the term “reproductive failure” to refer to women with UI, RM and RIF.

Uterine natural killer (uNK) cells are the most abundant immune cells found in placental bed of first trimester pregnancy, accounting for up to 70% of immune cells (Bulmer *et al*., 1991; King *et al*., 1991). It is widely recognized that uNK are important for the development of the placenta by facilitating trophoblast invasion and vascular remodelling (Moffett-King, 2002; Huhn *et al*., 2021). However, there is still uncertainty about how NK cells are associated with reproductive failure.

Initially, it was thought that uNK, similar to peripheral blood NK cells (pNK), exert cytotoxicity and have the potential to kill invading trophoblasts (Chao *et al*., 1995; Ruiz *et al*., 1996). However, this was quickly shown to be unlikely because uNK are less able to kill classical NK cell targets than pNK and have almost no ability to kill trophoblasts (King, Birkby and Loke, 1989).

Immunogenetic studies have revealed interesting discoveries about interactions between uNK receptors KIR, LILRB1 and CD94 with their ligands HLA-C, HLA-G and HLA-E, respectively, on extravillous trophoblast cells (EVT). The most compelling evidence comes from studies on KIR/HLA-C interaction. KIR expressed by uNK can confer either an activating or inhibitory signal, which in turn is dependent on the genes inherited by each individual. The genetic polymorphism of KIR can be simplified by considering two main KIR haplotypes, KIR A and KIR B. KIR A is comprised of mostly inhibitory genes such as KIR2DL2, KIR2DL3 and KIR2DL4 while KIR B is predominantly comprised of activating genes such as KIR2DS1 and KIR2DS2, although the inhibitory KIR2DL1 is also present in KIR B. The ligand for KIR is HLA-C molecule expressed by all EVT and can either be C1 or C2 epitope. The combination of maternal KIR and paternally derived HLA-C in fetus can be widely diverse, resulting in net activation or inhibition of NK cells (Moffett *et al*., 2016). In a study of women with RM and RIF undergoing assisted reproduction using their own or donor oocytes, women with a KIR AA genotype had lower livebirth rates compared to those with KIR AB or KIR BB genotypes, suggesting that reduced uNK activation is associated with negative reproductive outcomes (Alecsandru *et al*., 2020). The livebirth rate was also decreased as fetal HLA-C2 load is increased which has implications for future clinical utility in selection of fetus with HLA-C that is “compatible” with maternal KIR. (Alecsandru *et al*., 2020). However, this is still considered to be at the pre-clinical stage owing to several limitations of present evidence and practical considerations in the clinical setting (Moffett *et al*., 2016).

On the other hand, it has been proposed that increased uNK number or cytokine production could cause excessive premature angiogenesis which in turn cause harmful oxidative stress to the feto-placental unit (Chen *et al*., 2017). Research on uNK cytokine production has been inconclusive, with conflicting reports on which cytokine is excessive or deficient in reproductive failure (Woon *et al*., 2022). Potential reasons for this are the use of different assays to measure cytokine level, addition of other cytokines (like IL-2 or IL-15) into culture to recover immune cells which alters their cytokine production profile, and use of decidual samples collected after miscarriage potentially introducing the confounding factor of inflammatory processes that occur after a miscarriage (Moffett and Shreeve, 2022).

Since the landmark discovery of three uNK subsets by single-cell RNA sequencing (scRNAseq) (Vento-Tormo *et al*., 2018), it has become apparent that these subsets exhibit different phenotypes and function and thus are likely to have different roles in pregnancy (Huhn *et al*., 2021). A better characterization of the three uNK subsets was identified to be essential to the field (Huhn *et al*., 2021). We have recently characterized these uNK subsets throughout the normal human reproductive cycle (Whettlock *et al*., 2022) and have shown that all three subsets upregulate KIR and LILRB1 expression during first trimester. uNK also have the ability to increase cytokine production during the mid-secretory phase of the menstrual cycle, which aligns with period of embryo implantation. Further scRNAseq data have demonstrated that uNK1 frequency is lower and uNK3 higher in women with RM compared to pregnant controls (Guo *et al*., 2021; Wu *et al*., 2021).

However, these studies were performed in decidua collected after miscarriage which may have been affected by inflammatory processes thus not reflecting the immune environment before the miscarriage. Therefore, the question remains, how do uNK subsets behave in the endometrium of women with reproductive failure? Is one subset more affected than the others? Elucidating the answer to this may help us to focus research on a particular uNK subset to understand the pathophysiology behind reproductive failure. Therefore, we aim to evaluate differences in receptor expression, activation/cytokine production and proportion of matched pNK and uNK subsets in women with reproductive failure compared to controls and in women with ongoing compared to no ongoing pregnancy after in vitro fertilization (IVF).

## Materials and methods

### Primary tissue

Collection of human tissue was approved by Solihull Research Ethics Committee (study number: 22/WM/0041). Inclusion criteria for patient groups were as follows: Infertility defined as women below 40 years old with failure to conceive after regular unprotected intercourse for more than one year, RM defined as loss of two or more clinical pregnancies and RIF defined as absence of pregnancy after two or more fresh or frozen transfer of good quality embryos. Exclusion criteria were any other causes identified for the reproductive failure after routine investigations including uterine, tubal, ovarian, male factor, endocrinological, parental chromosomal abnormalities and haematological (e.g. thrombophilia and antiphospholipid syndrome) factors. Inclusion criteria for the control group were women below 40 years old undergoing coil insertion for contraception with a regular menstrual cycle between 25 to 35 days and no history of reproductive problems. Exclusion criterion was usage of any coil or hormonal contraception within one month prior to sampling. The research participants’ characteristics are summarised in Supplementary Table 1.

Matched peripheral blood and endometrial samples were obtained for 16 reproductive failure patients (9 UI, 4 RM, 3 RIF) and 11 controls during mid-luteal phase. Endometrial biopsies were obtained by Pipelle sampler, immediately suspended in RPMI and processed within 2 hours. For the control group, endometrial samples were obtained before coil insertion for contraception between day 18-23 and mid-luteal phase was confirmed by histological dating and serum progesterone level. For the patient group, samples were obtained seven to nine days after ovulation as determined by serial urine luteinizing hormone (LH) tests.

For extraction of peripheral blood lymphocytes (PBL), whole blood was layered onto Histopaque (Sigma Aldrich), centrifuged (700 x*g* 21°C, 20 minutes) and the enriched PBL were washed twice (500 x*g*, 10 minutes, 4°C) with Dulbecco’s Phosphate-Buffered Saline (Life Tech). For endometrial lymphocytes extraction, endometrial tissue was pressed through 100 µm cell strainer, pelleted (700 x*g*, 10 minutes, 4°C), resuspended in Dulbecco’s Phosphate-Buffered Saline supplemented by 10% Fetal Calf Serum (Sigma Aldrich), passed through 70µm strainer, and layered on Histopaque to isolate immune cells as above. 0.2 to 1 ×10^6^ endometrial lymphocytes and 1 ×10^6^ PBL were allocated per condition of either fresh stain or culture.

### Short-term culture for measurement of cytokine production

Both PBL and endometrial lymphocytes were suspended in RPMI supplemented with antibiotics, non-essential amino acids 10% fetal calf serum, 1 µM sodium pyruvate, 2.5 mM HEPES and 50 µM β-mercaptoethanol (all from Gibco) and divided into unstimulated and stimulated wells. Anti-CD107a BV605 (100 µl/ml; clone H4A3, Biolegend), Brefeldin (10µg/ml) and Monensin (2µM/ml) were added to all wells and Phorbol 12-myristate 13-acetate (PMA) (50 ng/ml) and ionomycin (1µg/ml) into the stimulated wells only. Cells were incubated for 4 hours at 37°C then harvested and stained.

### Flow cytometry

The following anti-human antibodies were used for surface staining: Anti-CD56 Brilliant Violet (BV) 650 (clone NCAM 16.2, BD Bioscience), anti-CD39 BV421 (clone A1, Biolegend), anti-CD3 BV711 (clone SK7, Biolegend), anti-CD103 BV785 (clone Ber-ACT8, Biolegend), anti-CD16 Alexa Fluor(AF)700 (clone 3G8, Biolegend), anti-CD9 phycoerythrin(PE)/Dazzle 594 (clone HI9a, Biolegend), anti-CD49a PE/Cy7 (clone TS2/7, Biolegend), anti-CD45 allophycocyanin (APC) (clone HI30, Biolegend), anti-CD94 PE (Clone HP-3D9, BD Bioscience), anti-CD158a/h (KIR2DL1/DS1) VioBright 515 (clone REA1010, Miltenyi Biotec), anti-CD158b (KIR2DL2/DL3) APC vio 770 (clone REA 1006, Miltenyi Biotec), CD85j (ILT2 or CD94) Peridinin chlorophyll protein (PerCP)-eFluor 710 (clone HP-F1, Thermo Fisher Scientific). The following anti-human antibodies were used for intracellular cytokine staining: anti-IL-8 PE (clone G265-8, BD Bioscience), anti-IFN-γ APCvio770 (clone REA600, Miltenyi Biotec), anti GM-CSF PERCP/Cyanine 5.5 (clone BVD2-21C11, Biolegend), anti-TNFα FITC (clone MAb11, Biolegend).

Cells were incubated with fixable viability dye (Live/Dead Fixable Aqua Dead Cell stain kit, LifeTech) and surface antibodies (15 minutes, 4°C). For intracellular staining, human FoxP3 buffer (BD Biosciences) was used according to manufacturer’s instructions for fixation and permeabilization before staining with intracellular antibodies (30 minutes, 4°C). Excess antibodies were washed off (5 minutes, 500 *xg*, 4°C) between each incubation and twice after the final incubation with intracellular antibodies.

### Statistical analysis

Data were acquired on an BD Fortessa and analysed using FlowJo (Tree Star, Ashalnd, OR). In order to ensure reproducibility of results, CS&T and application settings were used. Statistical analysis was performed using PRISM (GraphPad Software Inc.). Data were assessed for normality using Shapiro-Wilk tests to determine whether a parametric or a non-parametric statistical test was appropriate. The appropriate statistical test was used to compare patient and control groups as specified in figure legends. p<0.05 was considered significant.

## Results

### No difference in uNK or pNK frequencies in women with reproductive failure compared to controls

In women with recurrent miscarriage, assessment by scRNAseq on decidual samples after miscarriage found that uNK1 level is lower but uNK3 level is higher compared to controls, although these findings may represent inflammatory changes subsequent to fetal demise (Guo *et al*., 2021; Wu *et al*., 2021). Therefore, we evaluated proportion of uNK and its subsets from mid-luteal phase endometrial samples as total uNK, uNK1, 2 and 3 as proportion of CD45+ lymphocytes and the latter three as proportion of total uNK. NK cells were identified as live CD3-CD56+ lymphocytes, and CD49a was used to distinguish uterine from circulating NK cells. The uNK subsets were subsequently identified by CD103 and CD39 expression (Vento-Tormo *et al*., 2018) (Fig. 1A). pNK were identified by conventional gating strategy; CD3-CD56+ cells with gating by CD56 expression level and on CD16 to distinguish CD56^bright^ and CD56^dim^.

**Figure 1.**
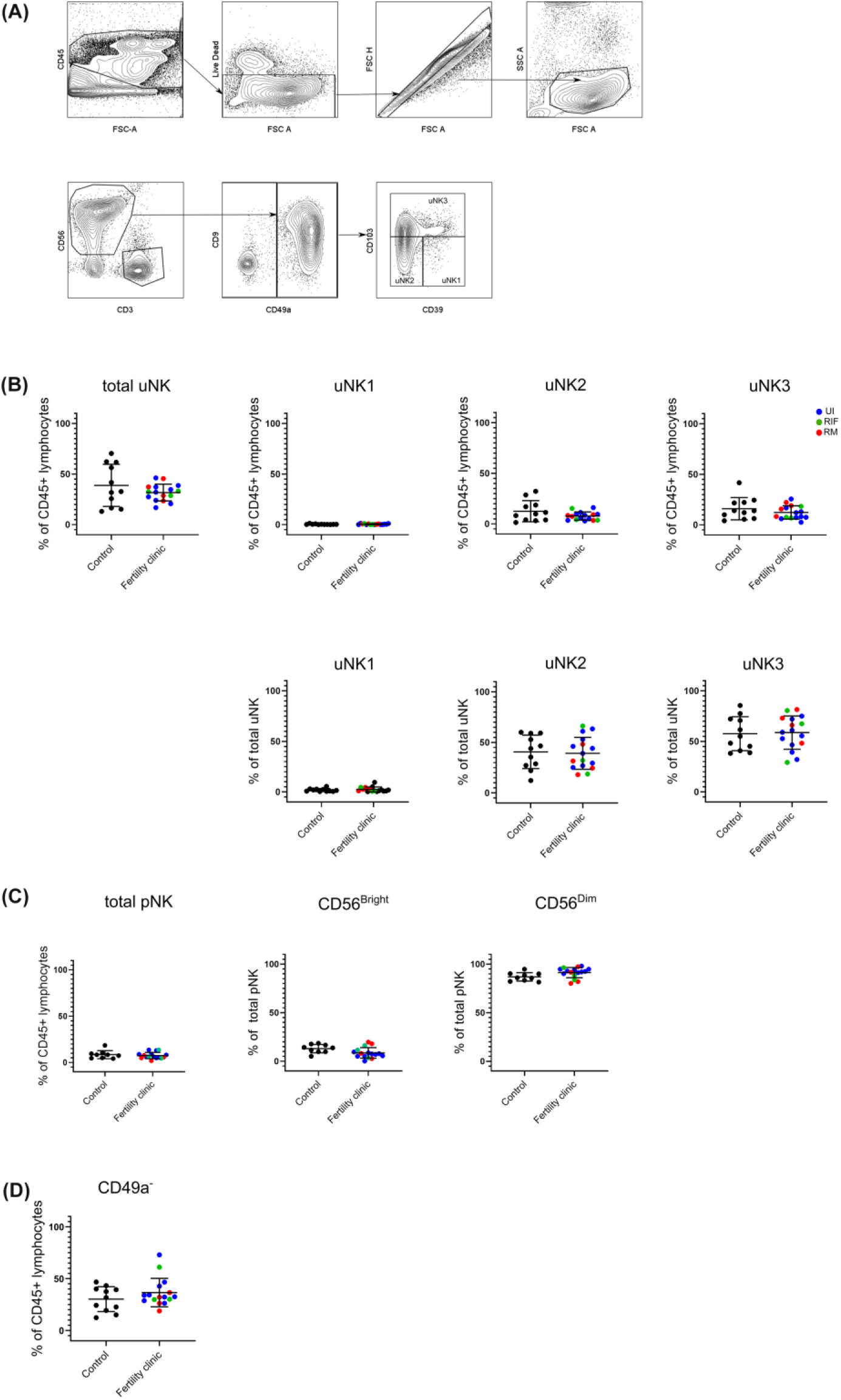
uNK and pNK proportion in unexplained subfertility, recurrent miscarriage and implantation failure compared to controls. **(A)** FACs gating strategy used to identify three uNK subsets (representative example shown) **(B)** Graphs of total uNK from CD45+ lymphocytes then frequency of each uNK subset (uNK1, -2, -3) as proportion of CD45+ lymphocytes and proportion of total uNK. **(C)** Graphs of total pNK from CD45+ lymphocytes then frequency of CD56^Bright^ and CD56^Dim^ as proportion of total pNK. **(D)** Graph of CD3-CD56+CD49a-lymphocytes as proportion of total uNK. Means and standard deviations are shown for controls (n=11) shown in black and patients from fertility clinic including unexplained infertility (UI; n=9) showed in blue, recurrent miscarriage (RM; n=34) shown in red and recurrent implantation failure (RIF; n=3) shown in green. Statistical testing was done using unpaired t-test for normal or Mann-Whitney U test for non-normal distribution.

In keeping with our previous finding at this stage in the menstrual cycle (Whettlock, 2022) uNK1 frequency was lower than that of uNK 2 and 3 in the endometrium in both control and UI/RM/RIF groups (Fig. 1B). However, there was no significant difference in frequency of total uNK or uNK subsets in reproductive failure compared to control group (Fig. 1B). Similarly, there was no significant difference in pNK (Fig. 1C). Additionally, no significant difference was detected even when each subgroup of reproductive failure was considered separately (data not shown).

In our previous meta-analysis, it was found that CD16+ cells, which are likely to represent CD56^dim^ pNK, are more frequency in the endometrium of women with recurrent miscarriage compared to controls (Woon *et al*., 2022). Therefore, we examined the frequency of CD3-CD56+CD49a-NK cells in the endometrium, since these more accurately represent pNK. Overall, no significant difference was detected in reproductive failure compared to control group (Fig 1D).

### KIR2DL1/S1 and LILRB1 expression are lower in women with reproductive failure

KIR, LILRB1 and CD94 are uNK receptors that interact with HLA-C, HLA-G and HLA-E on EVT respectively (Parham and Moffett, 2013). These receptors are present in all uNK subsets throughout the menstrual cycle and in all trimesters of pregnancy, albeit at different frequencies. Previously, we have shown that KIR2DL1/S1 and LILRB1 were upregulated in all three uNK subsets during first trimester, suggesting that these receptors are important for cross-talk with EVT at the time of placentation (Whettlock *et al*., 2022).

Here, we assessed KIR2DL1/S1, KIR2DL2/S2/L3, LILRB1 and CD94 expression on total uNK, uNK1, uNK2, uNK3 from mid-luteal phase endometrial samples and CD56^bright^ and CD56^dim^ pNK from matched peripheral blood samples in women with reproductive failure (RM, RIF or UI) compared to controls attending the clinic for contraceptive coil fitting. Representative histograms of staining from each receptor are shown in Fig. 2A.

**Figure 2.**
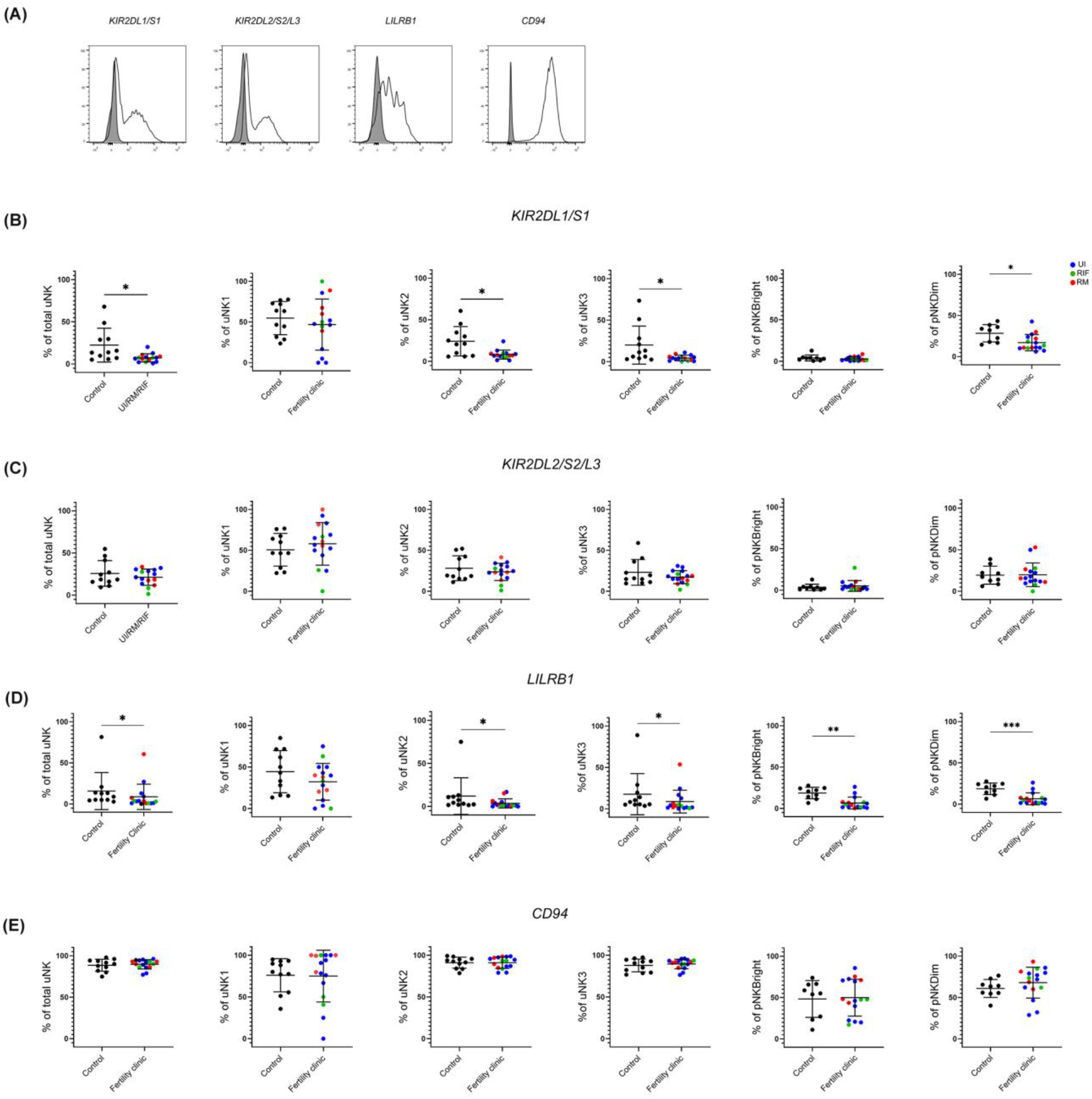
KIR2DL1/S1 and LILRB1 expression is lower in women with reproductive failure. **(A)** Uterine and peripheral NK cells taken from control and UI, RM and RIF groups were freshly stained for phenotypic markers. Representative staining of two samples from total uNK are shown in clear alongside fluorescence minus one control in grey. Graphs showing frequencies of KIR2DL1/S1 **(B)**, KIR2DL2/3 **(C)**, LILRB1 **(D)** and CD94 **(E)** on uNK and pNK. Means and standard deviations are shown for controls (n=11) shown in black and patients from fertility clinic including unexplained infertility (UI; n=9) showed in blue, recurrent miscarriage (RM; n=4) shown in red and recurrent implantation failure (RIF; n=3) shown in green. Statistical testing was done using unpaired t-test for normal or Mann-Whitney U test for non-normal distribution. * p<0.05, ** p<0.01, ***p<0.001.

KIR2DL1/S1 expression was significantly lower in total uNK in the reproductive failure group, with further investigation of expression in subsets revealing lower expression in all three uNK subsets, although this only reached statistical significance for uNK2 and 3 (Fig. 2B) In peripheral blood, KIR2DL1/S1 was also observed to be significantly lower in the reproductive failure group among CD56^dim^ but not CD56^bright^ cells (Fig. 2B). LILRB1 displayed a similar patten of expression, being significantly lower in total uNK and all three uNK subsets in the reproductive failure group, although this only reached statistical significance for uNK2 and 3 (Fig. 2D). In the blood, LILRB1 expression was lower in for the reproductive failure group for both CD56^dim^ and CD56^bright^ pNK (Fig. 2D). However, no trend was noted for KIR2DL2/S2/L3 (Figure 2C) or CD94 (Fig. 2E) in either uNK or pNK.

### uNK are less activated in women with reproductive failure

Degranulation, as measured by the internalisation of anti-CD107a is a good measure of uNK activation (Xiong *et al*., 2013; Kennedy *et al*., 2016) and using this measurement we have previously found that uNK are most active at the time of implantation during secretory phase and first trimester (Whettlock *et al*., 2022). CD107a trended upwards in secretory phase and was significantly higher in first trimester compared to third trimester in uNK2 and 3 after stimulation. We also found that TNFα, IFNγ and IL-8 production across all uNK subsets is highest after stimulation with PMA and ionomycin during secretory phase (Whettlock *et al*., 2022).

Here, we replicated the assessment for functional responses with and without stimulation on uNK and pNK from women with reproductive failure compared to controls (Fig. 3). Strikingly, we observed significantly lower degranulation by total uNK in women with reproductive failure, compared to controls, without stimulation (Fig. 3B). When uNK subsets were considered separately, the reduction was most pronounced on uNK3 although this did not reach statistical significance.

**Figure 3.**
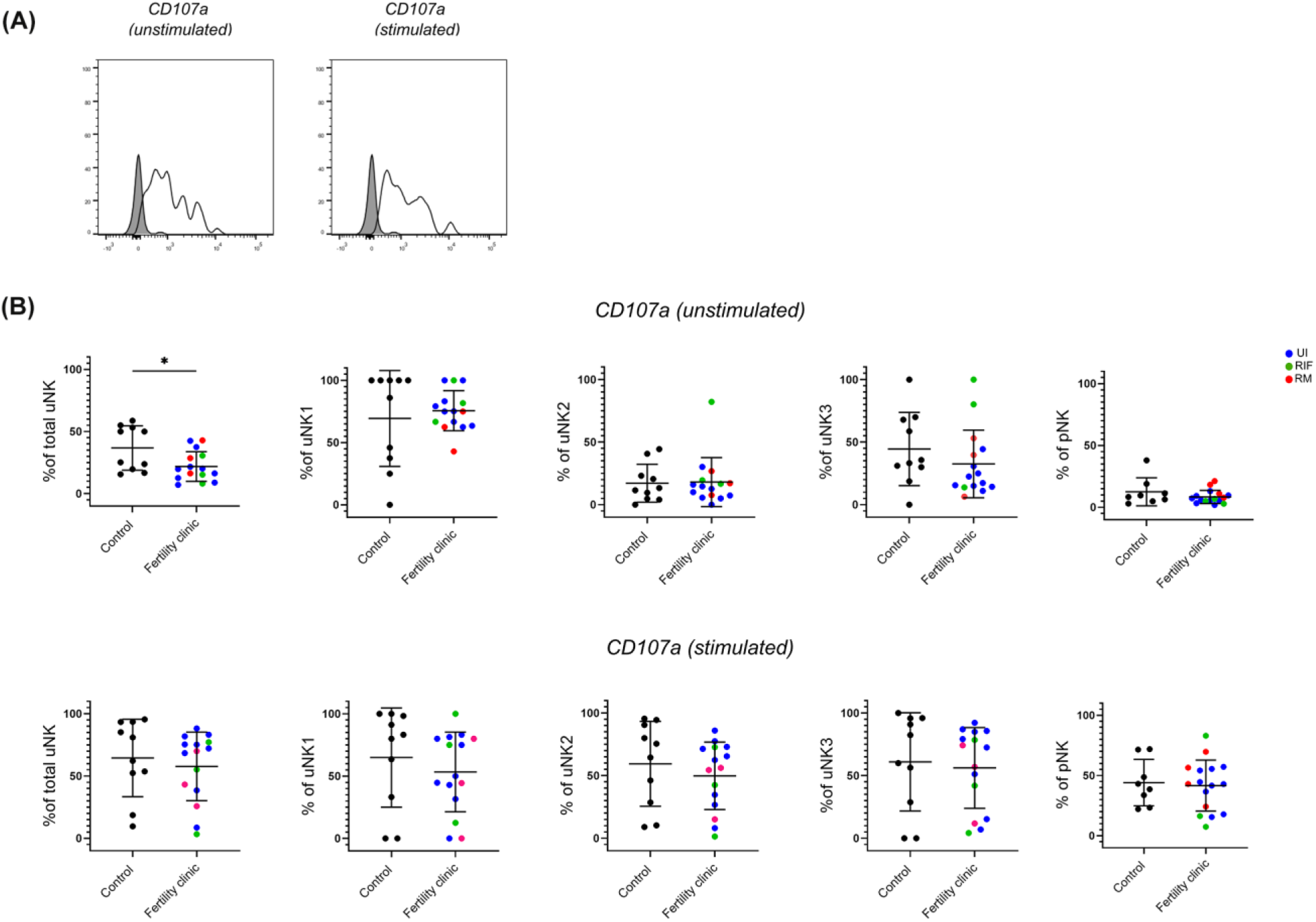
uNK is less active in women with reproductive failure. **(A)** Uterine and peripheral NK cells taken from control and UI, RM and RIF groups were cultured with or without PMA and ionomycin stimulation. Representative staining of one sample from total uNK is shown in clear alongside fluorescence minus one control in grey. Graphs showing frequencies of CD107a on unstimulated **(B)** and stimulated **(C)** uNK and pNK. Means and standard deviations are shown for controls (n=11) shown in black and patients from fertility clinic including unexplained infertility (UI; n=9) showed in blue, recurrent miscarriage (RM; n=3) shown in red and recurrent implantation failure (RIF; n=3) shown in green. Statistical testing was done using unpaired t-test for normal or Mann-Whitney U test for non-normal distribution. * p<0.05

However, no discernible difference was seen for degranulation by total uNK or uNK subsets with stimulation (Fig. 3C). Among all of the cytokines (TNFα, IFNγ, IL-8 and GM-CSF) that were assessed, TNFα emerged as the only one that was significantly lower in all three uNK subsets in the reproductive failure group with stimulation (Supp Fig. 1A) Notably, none of the functional changes in uNK were mirrored by pNK. (Fig. 3 and Supp, Fig. 1)

### No difference in phenotype or function of uNK in women with ongoing versus no ongoing pregnancy after IVF

Next, we prospectively followed up women from the reproductive failure group to obtain data on their pregnancy outcome after IVF. We compared women who had an ongoing pregnancy (OP) with women with no ongoing pregnancy (NOP). The latter group consisted of women who did not fall pregnant as well as women who fell pregnant but miscarried. Due to the small numbers from each subgroup, they were pooled into the NOP group. The same phenotypic and functional parameters as above were assessed.

No significant difference was seen in any of the KIR (Fig. 4A, 4B), LILRB1 (Fig. 4C) or CD94 (Fig. 4D) expression in either uNK or pNK. However, a trend of lower KIR expression was seen in total uNK and uNK subsets for the NOP group. On removal of the two datapoints of women who fell pregnant but miscarried from the NOP group, KIR2DL2/S2/L3 was significantly lower specifically in the uNK3 subgroup (data not shown). No other significant difference emerged from similar sensitivity analysis on the other receptors. In terms of functional activity, no significant difference was observed in CD107a (Fig. 4E, 4F) or any of the cytokines examined (Supp. Fig. 2).

**Figure 4.**
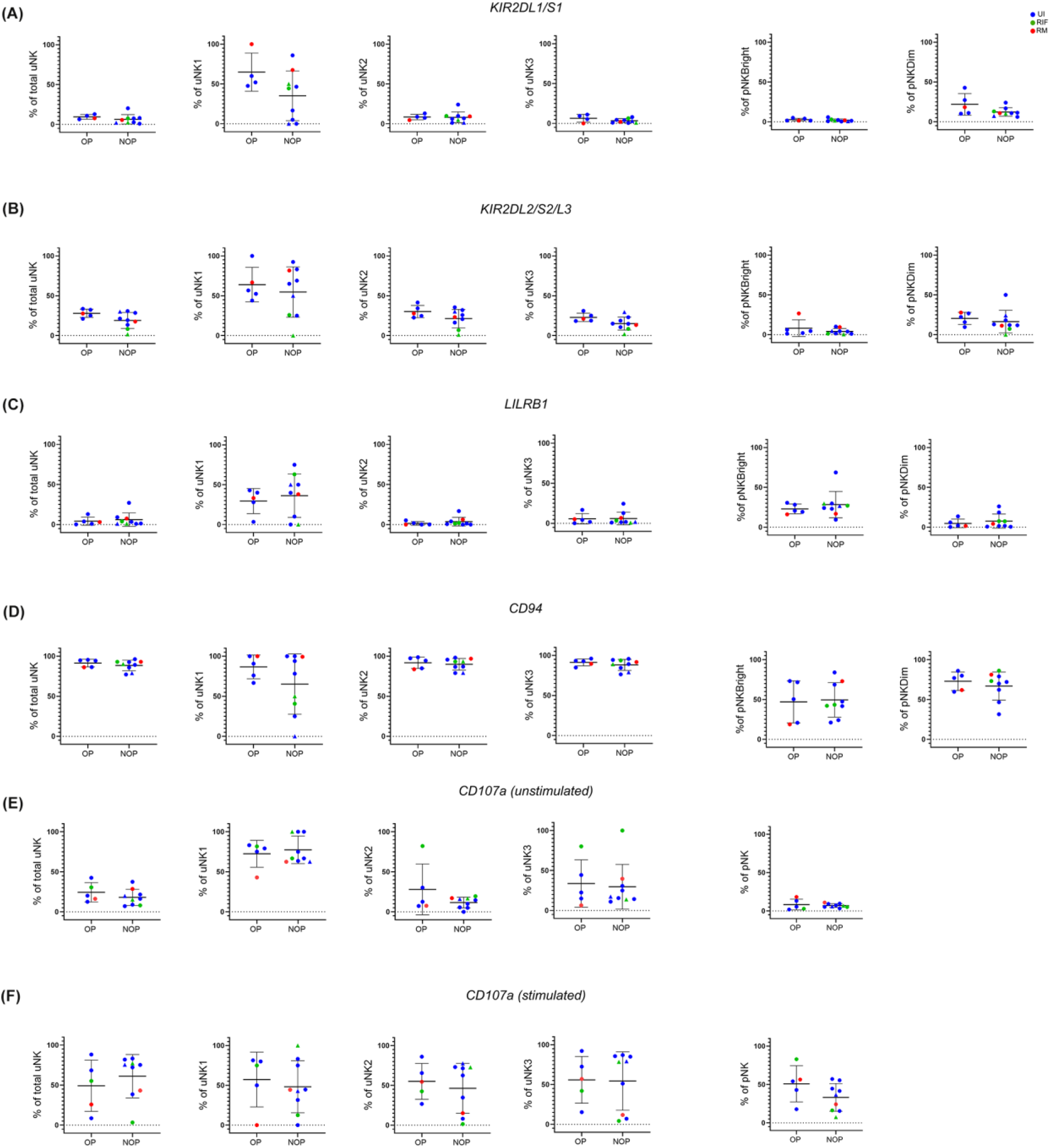
No difference in phenotype or degranulation in patients who had ongoing pregnancy compared to no ongoing pregnancy after IVF. Means and standard deviations are shown for KIR2DL1/S1 **(A)**, KIR2DL2/S2/L3 **(B)**, LILRB1 **(C)**, CD94 **(D)**, CD107a with **(E)** and without **(F)** stimulation. NOP group consist of n=7 who did not become (represented as dots) and n=2 who became pregnant but miscarried (represented as triangles). Unexplained infertility (UI) is showed in blue, recurrent miscarriage (RM) is shown in red and recurrent implantation failure (RIF) is shown in green. Statistical testing was done using unpaired t-test for normal or Mann-Whitney U test for non-normal distribution. * p<0.05

## Discussion

In this study, we have examined the frequency, phenotype and function of pNK and uNK subsets from timed endometrial samples obtained from mid-luteal phase in women with reproductive failure compared to controls. In contrast to previous studies of endometrial samples from UI, RM or RIF patients, which analyzed total CD3-CD56+ cells, here we also present data on the three uNK subsets individually.

Our previous systematic review showed significantly raised uNK in women with RM and RIF when considering total CD56+ NK cells in the endometrium during mid-luteal phase, but not in decidua collected after miscarriage (Woon *et al*., 2022). In women with RM, two recent studies using scRNAseq and mass cytometry demonstrated reduced uNK1 and raised uNK3 level in RM group (Guo *et al*., 2021; Wu *et al*., 2021), albeit the studies compared termination samples to those collected after miscarriage, so the results could be confounded by inflammatory changes occurring subsequent to fetal demise (Moffett, 2022). In comparison, we did not observe any difference in the frequency of either CD56+ uNK or uNK subsets. One possibility could be that our cohort included patients with a mixture of types of reproductive failure, although each subgroup did not show any difference when considered separately. Another possibility is that this small study was not sufficiently powered to detect a difference, although no emerging trend was apparent from our results. A third possibility is that the comparison is not perfect because the timing of samples for control group was done by serum progesterone and histological dating instead of urine LH tracking, which increases the variance in the control group, thus reducing the power to detect differences.

However, it should also not be discounted that the lack of difference is partially due to significantly lower level of uNK1 in the endometrium compared to decidua in first trimester, therefore no inherent difference in the frequency of endometrial NK cells may be present.

We found significantly lower expression of KIR2DL1/S1 in uNK subsets in the reproductive failure group compared to controls, and these findings were mirrored in pNK. Our finding in pNK is consistent with previous work which has reported a reduction in the expression of KIR2DL1 expression by pNK associated with reproductive failure (Ntrivalas *et al*., 2002; Yamada *et al*., 2004; Faridi *et al*., 2011; Kniotek *et al*., 2021), although we did not find a reduction in KIR2DL2 expression, which has also been reported (Ntrivalas *et al*., 2002). Studies on uNK KIR expression have thus far included samples retrieved after TOP stratified to high and low risk of resistance index on uterine artery doppler (Wallace *et al*., 2015) or post-surgical management of miscarriage (Wang *et al*., 2014), which reported reduction in KIR2DL1/S1 but no change in KIR2DL2/L3/S2 associated with worse pregnancy outcomes. Therefore our findings that uNK and pNK from UI, RM and RIF patients express lower levels of KIR2DL1/S1 are in line with previous work looking at other disorders of pregnancy.

Our finding that KIRD2L1/S1 expression was reduced on both uNK and pNK suggests that the deficiency in KIR expression is not tissue-specific, which in turn could reflect particular KIR genotypes associated with reproductive failure. Early immunogenetic approaches demonstrated that women with KIR AA genotype in combination with fetus carrying HLA-C2 epitope are at higher risk of disorders of placentation associated with pre-eclampsia, and identified KIR2DS1, specifically, as protective (Hiby 2004, 2010). Similarly, in IVF patients maternal KIR AA genotype is associated with higher miscarriage rate compared to KIR AB and BB, and this risk is augmented by presence of HLA-C2 on fetus (Alecsandru *et al*., 2014, 2020). Furthermore, *in vitro* studies found that stimulation of the activating KIR2DS1 and KIR2DS4 on uNK release GM-CSF that promoted trophoblast invasion (Xiong 2013, Kennedy 2016). Collectively, these findings suggest that interaction with EVT that results in excessive inhibition of NK cells result in adverse pregnancy outcome, although some other studies have suggested that RM is associated with excess of NK cell activation (Flores *et al*., 2007; Faridi *et al*., 2009; Morin *et al*., 2016). A limitation of our approach is that the antibody used does not differentiate activating KIR2DS1 from inhibitory KIR2DL1: approaches that can achieve this may help to clarify whether reproductive failure is associated with uNK activation or inhibition via KIR.

We also observed reduced LILRB1 expression in both uNK and pNK from reproductive failure patients. Although LILRB1 ligation inhibits pNK, recent research has found that ligation of LILRB1on uNK by HLA-G promotes growth factor secretion by uNK which in turn promotes development of the fetus through the placenta (Fu *et al*., 2017). A subset of uNK which express NKG2C and LILRB1 has been reported in multiparous women (Gamliel *et al*., 2018). It has been suggested that these cells could account for the greater likelihood of success in second and subsequent pregnancies (Goldman-Wohl *et al*., 2019) and if so, this would be consistent with a role for LILRB1 in optimizing placental development. For women with reproductive failure, studies on LILRB1 have mostly concentrated on pNK in RM patients (Kniotek *et al*., 2021; Habets *et al*., 2022) except for one study on uNK using TOP samples stratified by uterine artery resistance (Wallace *et al*., 2014). On balance, and in line with our findings, these studies reported reduced expression of LILRB1 in women with high risk of reproductive problems.

We found significantly lower degranulation by total uNK in the reproductive failure group, although no one subset accounted for this reduction, and this finding lends weight to the hypothesis that underactivation of uNK is associated with reproductive failure. Notably, this finding was specific to uNK as there was no evidence of reduced degranulation in pNK from reproductive failure patients. Similarly, the only cytokine that was altered between controls and reproductive failure patients was TNFα, whose production after stimulation was reduced specifically in all uNK subsets, but not in pNK, from reproductive failure patients. This finding was in line with an earlier report that uNK from mid-luteal biopsies display reduced production of TNFα in women with RM Fukui *et al*., (2017). On the other hand, there are reports that RM is associated with increased uNK production of TNFα and IFN-γ, although a major limitation is that all of these studies were performed on decidua collected after miscarriage hence associated with the confounding factor of inflammatory processes occurring after a miscarriage. (Dong *et al*., 2017; Liu *et al*., 2019, 2020; Fonseca *et al*., 2020; Takeyama *et al*., 2021). There is some evidence that TNFα inhibits EVT invasion by mechanisms such as trophoblast apoptosis, inhibition of proliferation and reduced matrix metalloprotease production (Lash *et al*., 2006; Otun *et al*., 2011). However, more recently it has been suggested that implantation is a pro-inflammatory process, in which case TNFα production might be expected to be beneficial (Wang *et al*., 2020). In support of this theory, our previous results demonstrated that TNFα and IL-8 production by uNK peak in secretory phase, potentially indicating a role for these cytokines during the window of implantation (Whettlock *et al*., 2022). Here, we have shown deficiencies of uNK activation and TNFα production during mid-luteal phase in the reproductive failure group which supports the hypothesis that pro-inflammatory cytokines are important during the implantation window.

Taken together, our findings suggest global reduction of NK cell receptors important for cross-talk with HLA molecules on EVT, which may be genetically mediated, leads to reduced activation specifically in uNK, which in turn impairs implantation and placental development. However, our finding that expression of KIR and LILRB1, and uNK degranulation, is not predictive of successful pregnancy outcome sounds a note of caution about this interpretation, since it could suggest that the phenotypic and functional differences we observe are an effect, rather than a cause, of repeated failures of implantation. The insight gained from our findings will aid the direction of future research on uNK subsets in women with reproductive problems. Further larger cohorts should be recruited for studies on phenotype and function of uNK subsets, with a focus on KIR genotype and phenotype as well as uNK activation.

## Data availability

Raw data and analysis files have been deposited at the Open Science Framework, doi: 10.17605/OSF.IO/Y2PCD

## Author Contributions

EVW, MJ and VM designed the study. EVW, DN, KM and JNT participated in recruitment of patients and collection of clinical samples. PB performed histological dating. EVW and AOC carried out laboratory experiments. EVW and VM analysed results and wrote the manuscript. All authors contributed to editing the manuscript.

## Acknowledgments

We would like to thank all the people from Fertility Centre, Chelsea and Westminster Hospital and John Hunter Clinic (London, UK) who contributed samples to this study, as well as the medical, nursing and embryology staff who helped in the collection of samples. We would also like to thank Emily Whettlock for her contribution in optimizing the original flow cytometry panel.

