## Supplementary figures for "Uterine NK cells underexpress receptors recognizing HLA-C2 and HLA-G in reproductive failure"

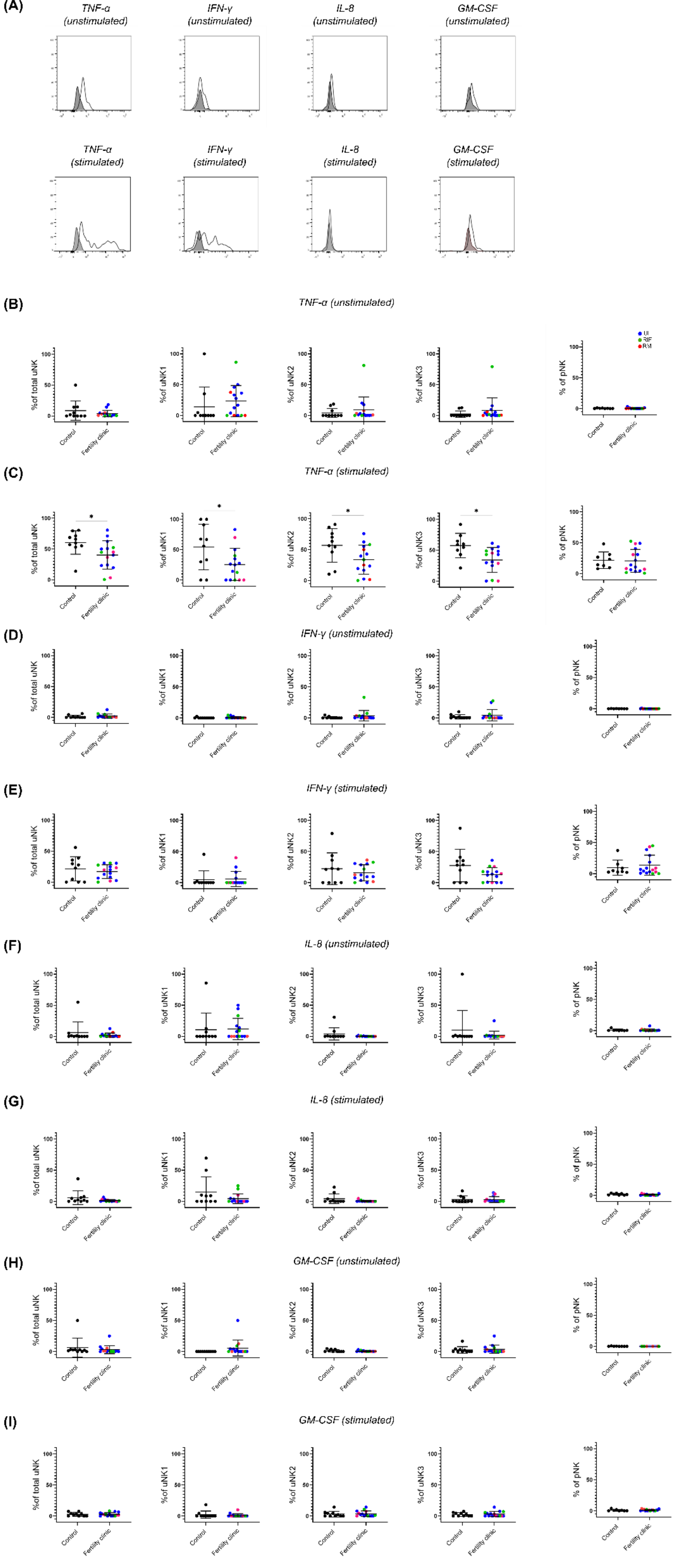

(A)
TNF-α (unstimulated)
IFN-γ (unstimulated)
IL-8 (unstimulated)
GM-CSF (unstimulated)
TNF-α (stimulated)
IFN-γ (stimulated)
IL-8 (stimulated)
GM-CSF (stimulated)
(B)
TNF-α (unstimulated)
(C)
TNF-α (stimulated)
(D)
IFN-γ (unstimulated)
(E)
IFN-γ (stimulated)
(F)
IL-8 (unstimulated)
(G)
IL-8 (stimulated)
(H)
GM-CSF (unstimulated)
(I)
GM-CSF (stimulated)
%of total uNK
%of uNK1
%of uNK2
%of uNK3
% of pNK
Control
Fertility clinic
0
50
100
*

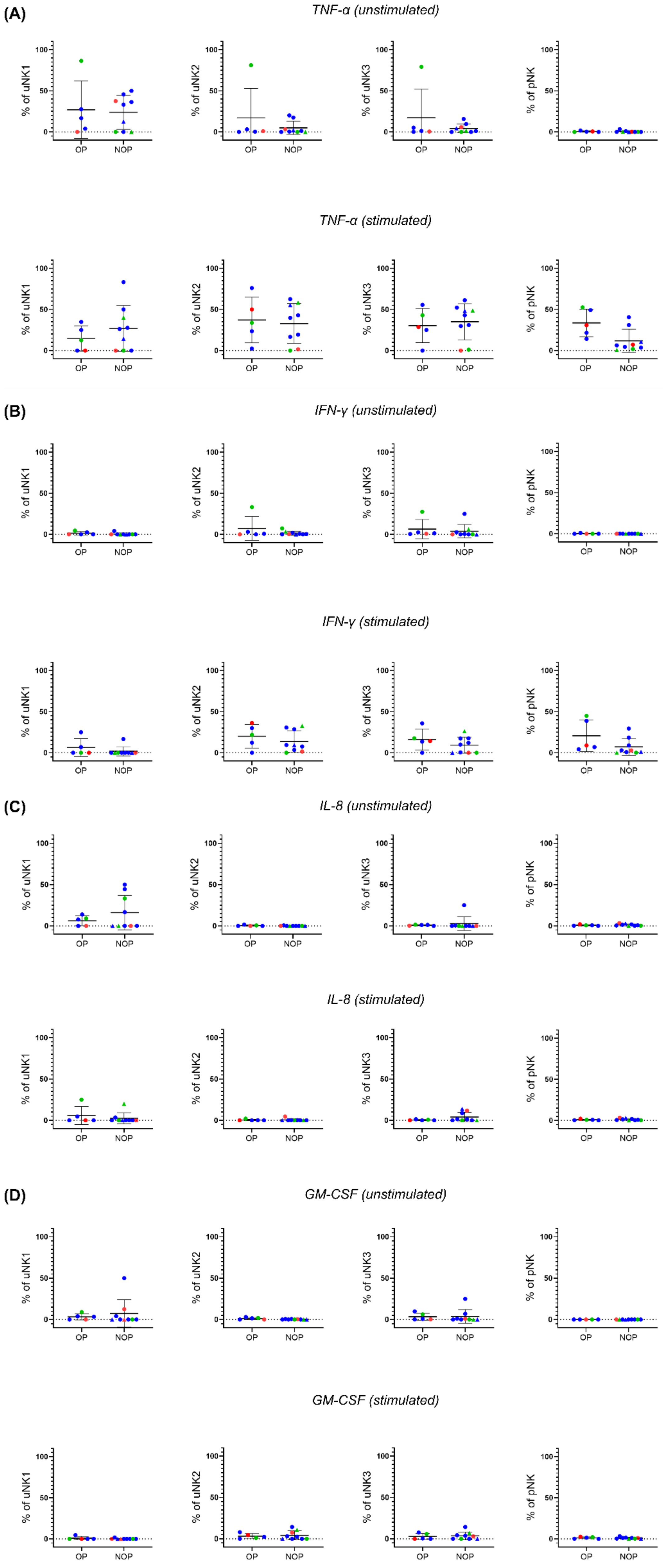

(A)
TNF-α (unstimulated)
TNF-α (stimulated)
(B)
IFN-γ (unstimulated)
IFN-γ (stimulated)
(C)
IL-8 (unstimulated)
IL-8 (stimulated)
(D)
GM-CSF (unstimulated)
GM-CSF (stimulated)
% of uNK1
% of uNK2
% of uNK3
% of pNK
OP
NOP

| Participant ID | Condition | Day of cycle | Length of cycle | Days post LH surge | Age | Gravida | Parity | Previous pregnancies | Previous IVF cycles | Pregnancy outcome | Serum progesterone | Assessment |
|---|---|---|---|---|---|---|---|---|---|---|---|---|
| 1 | UI | 22 | 28 | 9 | 32 | 0 | 0 | 0 | 1 | Pregnant | 17.9 | Phenotype, function, |
| 2 | UI | 20 | 28 | 7 | 39 | 0 | 0 | 0 | 0 | Not pregnant x2 | 51.7 | Phenotype, function, |
| 3 | UI | 22 | 28 | 9 | 34 | 0 | 0 | 0 | 0 | Pregnant | NI | Phenotype, function, |
| 4 | UI | 22 | 33 | 7 | 36 | 0 | 0 | 0 | 0 | Not pregnant | 36.2 | Phenotype, function, |
| 5 | UI | 23 | 33 | 7 | 36 | 0 | 0 | 0 | 0 | Not pregnant x2 | NI | Phenotype, function, |
| 6 | UI | 20 | 28-32 | 8 | 35 | 0 | 0 | 0 | 1 | Miscarriage | NI | Phenotype, function, |
| 7 | UI | 22 | 28 | 7 | 38 | 0 | 0 | 0 | 0 | Pregnant | 33.5 | Phenotype, function, |
| 8 | UI | 20 | 28 | 7 | 36 | 0 | 0 | 0 | 1 | Not pregnant | 35.7 | Phenotype, function, |
| 9 | UI | 21 | 28 | 7 | 35 | 0 | 0 | 0 | 0 | Not pregnant | NI | Phenotype, function, |
| 10 | RM | 20 | 28-30 | 7 | 39 | 3 | 0 | 3 miscarriages | 0 | FAE | NI | Phenotype, function for PBL only |
| 11 | RM | 22 | 28 | 9 | 34 | 3 | 0 | x3 miscarriages | x2 IVF - 1 no pregnancy, 1 miscarriage | Not pregnant | 31.8 | Phenotype, function |
| 12 | RM | 20 | 25-32 | 9 | 32 | 2 | 0 | x2 miscarriages | 0 | No outcome | NI | Phenotype, function |
| 13 | RM | 20 | 28 | 7 | 36 | 5 | 2 | x4 miscarriages | 0 | Pregnant | 39.7 | Phenotype, function |
| 14 | RIF | 25 | 30 | 7 | 39 | 1 | 1 | X1 livebirth with donor sperm | x3 failed implantation after IVF and ICSI | Pregnant | 39.5 | Phenotype, function |
| 15 | RIF+UI | 17 | 28 | 7 | 33 | 0 | 0 | 0 | x1IVF, x1 ICSI | Not pregnant | 42.1 | Phenotype, function |
| 16 | RIF+RM | 21 | 30 | 7 | 35 | 0 | 0 | x2 miscarriages | x3 IVF | Miscarriage | NI | Phenotype, function |
| 17 | Control | 20 | 28-30 | NI | 27 | 0 | 0 | N/A | N/A | N/A | 23 | Phenotype, function |
| 18 | Control | 19 | 28 | NI | 30 | 0 | 0 | N/A | N/A | N/A | 51 | Phenotype, function |
| 19 | Control | 21 | 28 | NI | 40 | 4 | 3 | 3 livebirths, 1 miscarriage | N/A | N/A | 28 | Phenotype, function |
| 20 | Control | 18 | 28 | NI | 25 | 0 | 0 | N/A | N/A | N/A | NI | Phenotype, function |
| 21 | Control | 19 | 28-30 | NI | 33 | 1 | 1 | 1 livebirth | N/A | N/A | 22 | Phenotype, function |
| 22 | Control | 17 | 28-30 | NI | 32 | 0 | 0 | N/A | N/A | N/A | 13 | Phenotype, function |
| 23 | Control | 21 | 28 | NI | 39 | 3 | 2 | 2 livebirths | N/A | N/A | NI | Phenotype, function |
| 24 | Control | 19 | 28 | NI | 27 | 0 | 0 | N/A | N/A | N/A | 24.2 | Phenotype, function |
| 25 | Control | 19 | 28 | NI | 31 | 0 | 0 | N/A | N/A | N/A | NI | Phenotype and function for EL only |
| 26 | Control | 19 | 30 | NI | 35 | 3 | 3 | 3 livebirths | N/A | N/A | NI | Phenotype and function for EL only |
| 27 | Control | 24 | 33 | NI | 25 | 0 | 0 | N/A | N/A | N/A | 8.1 | Phenotype, function |
